# Improving the accessibility and transferability of machine learning algorithms for identification of animals in camera trap images: MLWIC2

**DOI:** 10.1101/2020.03.18.997700

**Authors:** Michael A. Tabak, Mohammad S. Norouzzadeh, David W. Wolfson, Erica J. Newton, Raoul K. Boughton, Jacob S. Ivan, Eric A. Odell, Eric S. Newkirk, Reesa Y. Conrey, Jennifer L. Stenglein, Fabiola Iannarilli, John Erb, Ryan K. Brook, Amy J. Davis, Jesse S. Lewis, Daniel P. Walsh, James C. Beasley, Kurt C. VerCauteren, Jeff Clune, Ryan S. Miller

## Abstract

1. Motion-activated wildlife cameras (or “camera traps”) are frequently used to remotely and non-invasively observe animals. The vast number of images collected from camera trap projects have prompted some biologists to employ machine learning algorithms to automatically recognize species in these images, or at least filter-out images that do not contain animals. These approaches are often limited by model transferability, as a model trained to recognize species from one location might not work as well for the same species in different locations. Furthermore, these methods often require advanced computational skills, making them inaccessible to many biologists.
2. We used 3 million camera trap images from 18 studies in 10 states across the United States of America to train two deep neural networks, one that recognizes 58 species, the “species model,” and one that determines if an image is empty or if it contains an animal, the “empty-animal model.”
3. Our species model and empty-animal model had accuracies of 96.8% and 97.3%, respectively. Furthermore, the models performed well on some out-of-sample datasets, as the species model had 91% accuracy on species from Canada (accuracy range 36-91% across all out-of-sample datasets) and the empty-animal model achieved an accuracy of 91-94% on out-of-sample datasets from different continents.
4. Our software addresses some of the limitations of using machine learning to classify images from camera traps. By including many species from several locations, our species model is potentially applicable to many camera trap studies in North America. We also found that our empty-animal model can facilitate removal of images without animals globally. We provide the trained models in an R package (mlwic2: Machine Learning for Wildlife Image Classification in R), which contains Shiny Applications that allow scientists with minimal programming experience to use trained models and train new models in six neural network architectures with varying depths.

## 1 Introduction

Motion-activated wildlife cameras (or “camera traps”) are frequently used to remotely observe wild animals, but images from camera traps must be classified to extract their biological data (O’Connell, Nichols, & Karanth, 2011). Manually classifying camera trap images is an encumbrance that has prompted scientists to use machine learning to automatically classify images (Norouzzadeh et al., 2018; Willi et al., 2019), but this approach has limitations.

We address two major limitations of using machine learning to automatically classify animals in camera trap images. First, machine learning models trained to recognize species from one location and in one camera trap setup might perform poorly when applied to images from camera traps in different conditions. This “transferability problem” is thought to arise because different locations have different backgrounds (the part of the picture that is not the animal) and most models evaluate the entire image, including the background (Beery, Morris, & Yang, 2019; Miao et al., 2019; Norouzzadeh et al., 2019; Terry, Roy, & August, 2020; Wei, Luo, Ran, & Li, 2020). By including images from 18 different studies in North America, our objective was to train models with more variation in the backgrounds associated with each species. Furthermore, by training an additional model that distinguishes between images with and without animals, we provide an option that could be broadly applicable to camera trap studies worldwide. Second, the use of machine learning in camera trap analysis is often limited to computer scientists, yet the need for image processing exceeds the availability of computer scientists in wildlife research. To facilitate the use of these models by biologists with minimal programming experience, Machine Learning for Wildlife Image Classification (mlwic2) includes an option to train and use models in user-friendly Shiny Applications (Chang, Cheng, Alaire, Xie, & McPherson, 2019), allowing users to point-and-click instead of using a command line. This facilitates easier site-specific model training when our models do not perform to expectations.

## 2 Materials and Methods

### 2.1 Camera trap images

Images were collected from 18 studies using camera traps in 10 states in the United States of America (California, Colorado, Florida, Idaho, Minnesota, Montana, South Carolina, Texas, Washington, and Wisconsin; Appendix S1). Images were either classified by a single wildlife expert or classified independently by two biologists, with discrepancies settled by a third. An image was classified as containing an animal if it contained any part of an animal. Our initial dataset included 6.3 million images but was unbalanced with most images from a few species (e.g., 51% of all images were Bos taurus). We rebalanced the number of images by species and site to ensure that no one species or site dominated the training process. Previous work suggested that training a model with 100,000 images per species produces good performance (Tabak et al., 2019); therefore, we limited the number of images for a single species from one location to 100,000. When > 100,000 images for a single species existed at one location, we randomly selected 100,000 of these images to include in the training/testing dataset. After rebalancing the data, we had a total of 2.98 million images; 90% were randomly selected for training, while 10% were used for testing. Images used in this study were either already a part of or were added to the North American Camera Trap Images dataset (lila.science/datasets/nacti; Tabak et al., 2019). Images from Canada were not used for training but were used to evaluate model transferability as an out-of-sample dataset.

### 2.2 Training models

We trained deep convolutional neural networks using the ResNet-18 architecture (He, Zhang, Ren, & Sun, 2016) in the Tensorflow framework (Adabi et al., 2016) on a high performance computing cluster, “Teton” (Advanced Research Computing Center, 2018). Models were trained for 55 epochs, with a ReLU activation function at every hidden layer and a softmax function in the output layer, mini-batch stochastic gradient descent with a momentum hyperparameter of 0.9 (Goodfellow, Bengio, & Courville, 2016), a batch size of 256 images, and learning rates and weight decays that varied by epoch number (described in Appendix S2). We trained a species model, which contained classes for 58 species or groups of species and one class for empty images (Table 1). We also trained an empty-animal model that contained only two classes, one for images containing an animal, and the other for images without animals.

**Table 1:**
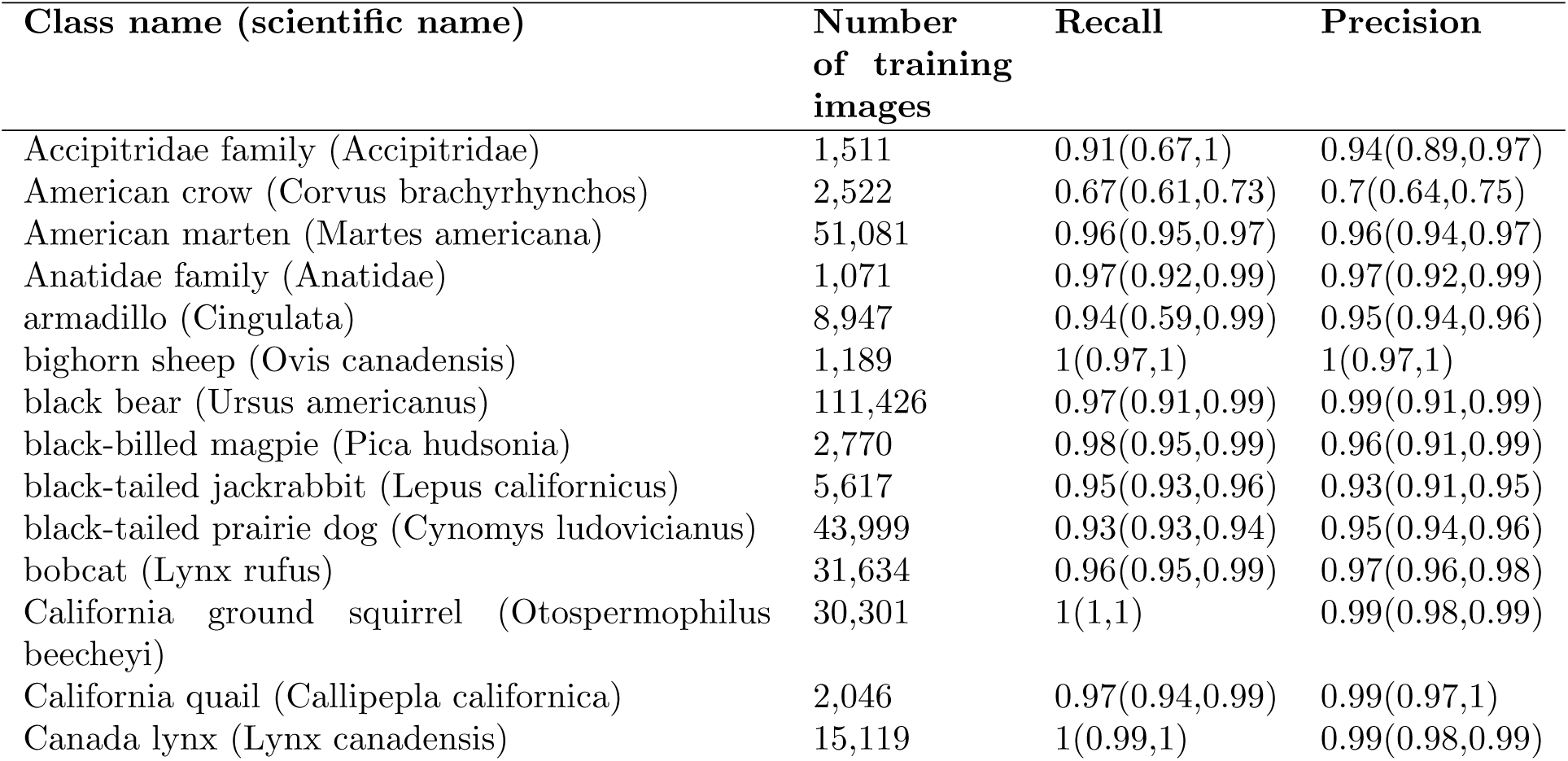
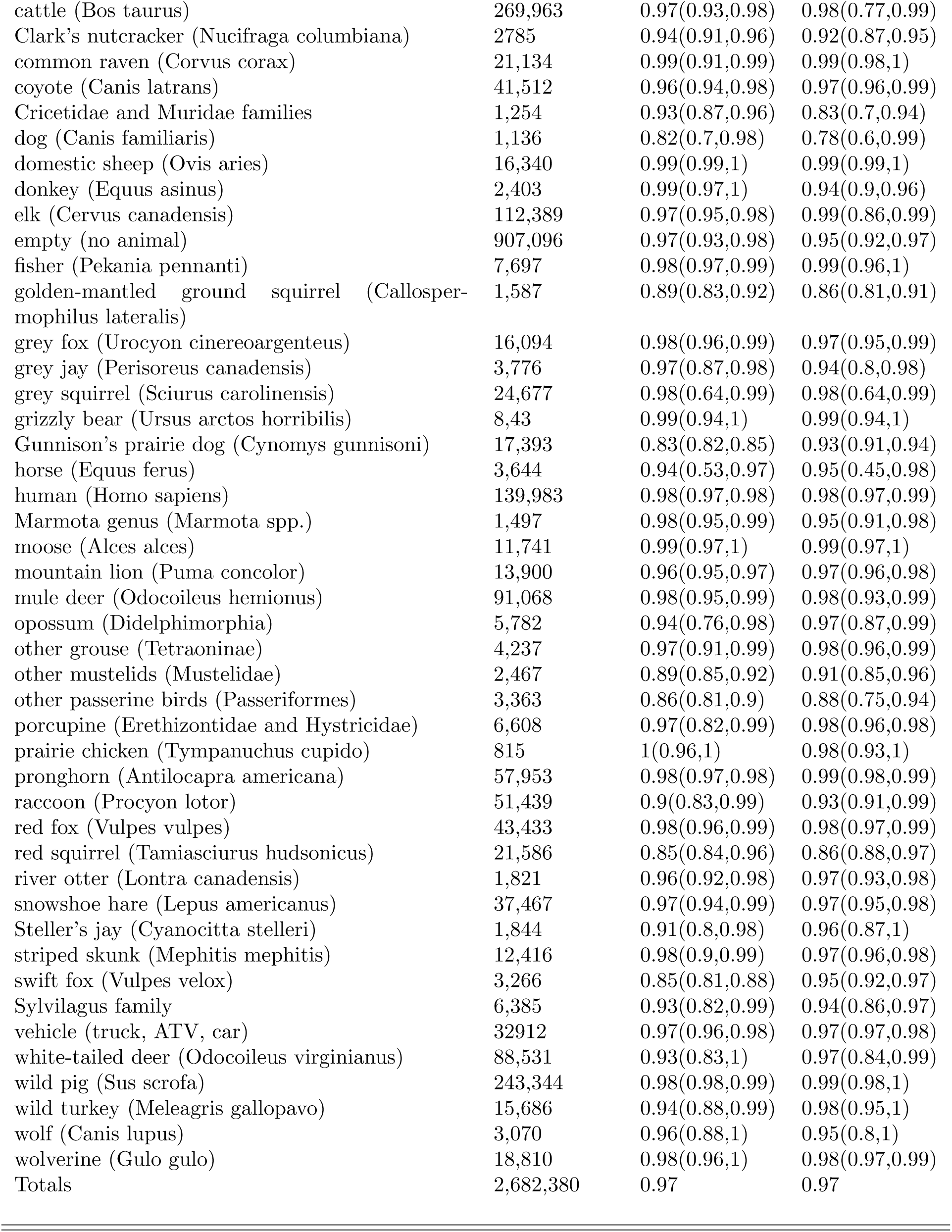
Mean recall and precision rates (along with 95% confidence intervals) for predicting species using the species model on the 10% of images that were withheld from training.

### 2.3 Model validation and transferability

We first evaluated our trained models by applying them to predicting species in the 10% of images that were withheld from training. Models were evaluated for each species using the recall, top-5 recall, and precision, which are values summarizing the number of true positives (TPs), false positives (FPs), and false negatives (FNs):

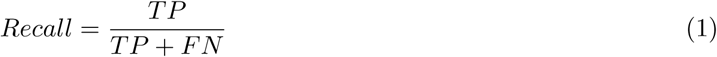

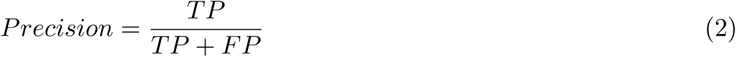

As recall is the proportion of images of each species that were correctly classified, top-5 recall is the proportion of images for each species in which one of the model’s top five guesses is the correct species. We also calculated confidence intervals for recall and precision rates (Appendix S3). To evaluate transferability of the model, we conducted out-of-sample validation by applying our trained models to images from locations where the model was not trained. We evaluated the species model using four out-of-sample datasets from North America: the Caltech Camera Traps dataset (Beery, Van Horn, & Perona, 2018), the ENA24-detection dataset (Yousif, Kays, & He, 2019), the Saskatchewan, Canada dataset from this study, and the Missouri Camera Traps dataset (Zhang, He, Cao, & Cao, 2016). The empty-animal model was tested using the Wellington Camera Traps dataset from New Zealand (Anton, Hartley, Geldenhuis, & Wittmer, 2018), the Snapshot Serengeti dataset from Tanzania (Swanson et al., 2015), and the Snapshot Karoo dataset from South Africa (http://lila.science/datasets/snapshot-karoo).

### 2.4 R package development

mlwic2 was developed using the R packages Shiny (Chang et al., 2019) and ShinyFiles (Pedersen, Nijs, Schaffner, & Nantz, 2019) so the user can choose to either use a programming console or a graphical user interface. Users can navigate to locations on their computer using a browser window instead of specifying paths. The package can classify images at a rate of 2,000 images per minute on a laptop with 16 gigabytes of random-access memory. mlwic2 will optionally write the top guess from each model and confidence associated with these guesses to the metadata of the original image file. The function `write_metadata` uses Exiftool (Harvey, 2016) to accomplish this. In addition, if scientists have labeled images, mlwic2 has a Shiny app that allows users to train a new model to recognize species using one of six different convolutional neural network architectures (AlexNet, DenseNet, GoogLeNet, NiN, ResNet, and VGG) with different numbers of layers. Note that the time required to train a model depends on the number of images used for training and computing resources; operating mlwic2 on a high-performance computing cluster requires programming experience.

## 3 Results

When we evaluated our models on the withheld images (within sample validation), we found an overall accuracy of 96.8% for the species model and 97.3% for the empty-animal model. Several species (six of 11) had recall of > 95% with fewer than 2,000 images used for training (Table 1; Fig.1). A confusion matrix (Appendix S4) depicts how all images of each species were classified by the species model. When evaluated on out-of-sample images, the species model accuracy ranged from 36.3% to 91.3% (Table 2), with top-5 accuracy ranging from 65.2% to 93.8% (Fig. 2), and the empty-animal model accuracy ranged from 90.6% to 94.1% (Table 2).

**Figure 1:**
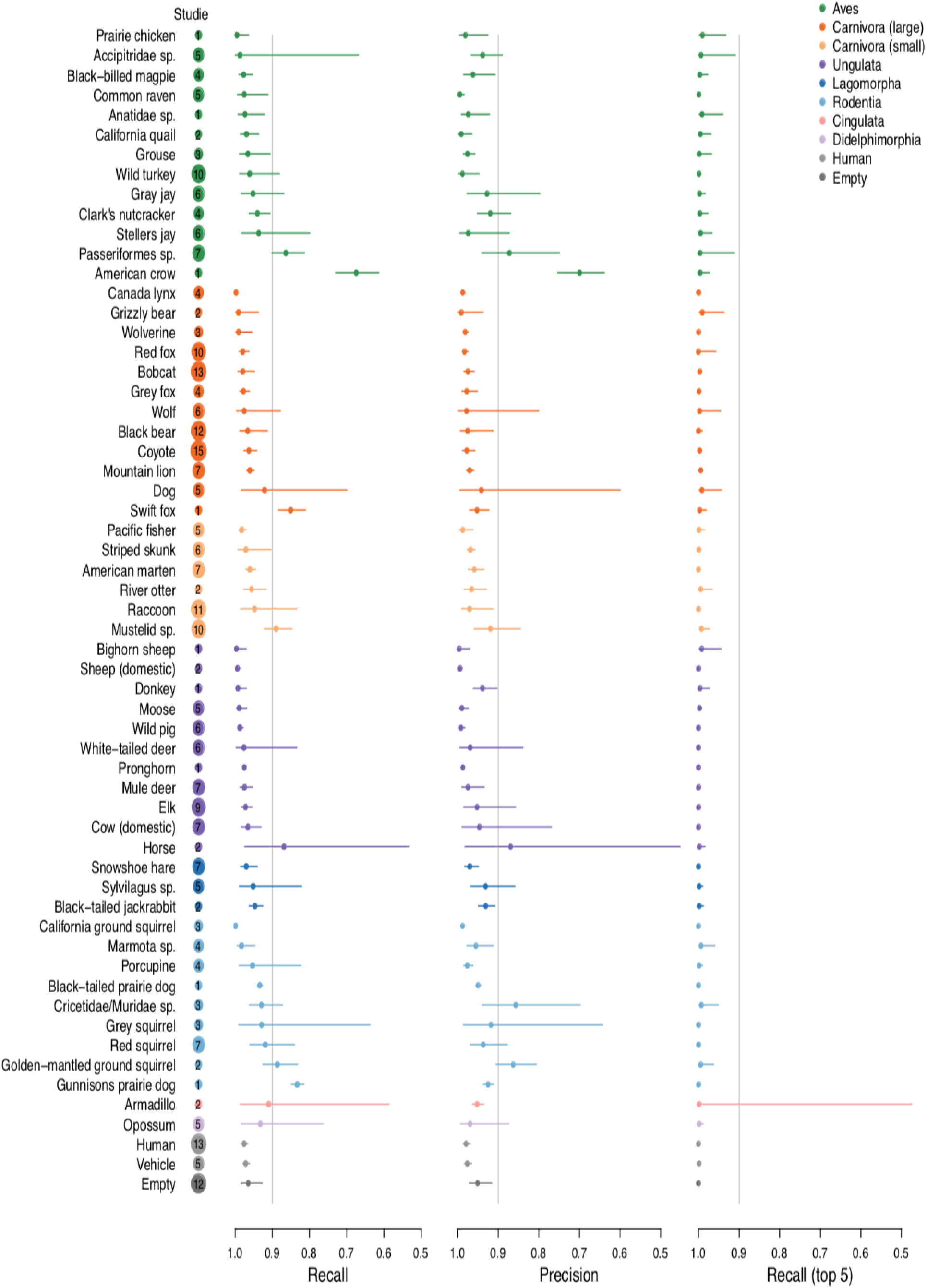
Within sample validation of the species model revealed high recall and precision for most species. Median values across datasets are presented along with 95% confidence intervals. The number of datasets for each species is included in the circle next to the species name (circle sizes are proportional to the number of datasets containing each species).

**Table 2:** Out-of-sample validation results. All out-of-sample images are available from lila.science/datasets.

| Dataset | Number of images tested | Model tested | Accuracy | Top-5 accuracy* |
| --- | --- | --- | --- | --- |
| Snapshot Karoo (South Africa) | 38,101 | empty-animal | 0.906 |  |
| Snapshot Serengeti (Tanzania) | 104,651 | empty-animal | 0.941 |  |
| Wellington (New Zealand) | 266,966 | empty-animal | 0.939 |  |
| Caltech Camera Traps (USA) | 218,147 | species | 0.562 | 0.744 |
| ENA24-Detection (USA) | 5,285 | species | 0.507 | 0.649 |
| Missouri Camera Traps (USA) | 5,008 | species | 0.362 | 0.652 |
| Saskatchewan (Canada) | 5,200 | species | 0.913 | 0.938 |

**Figure 2:**
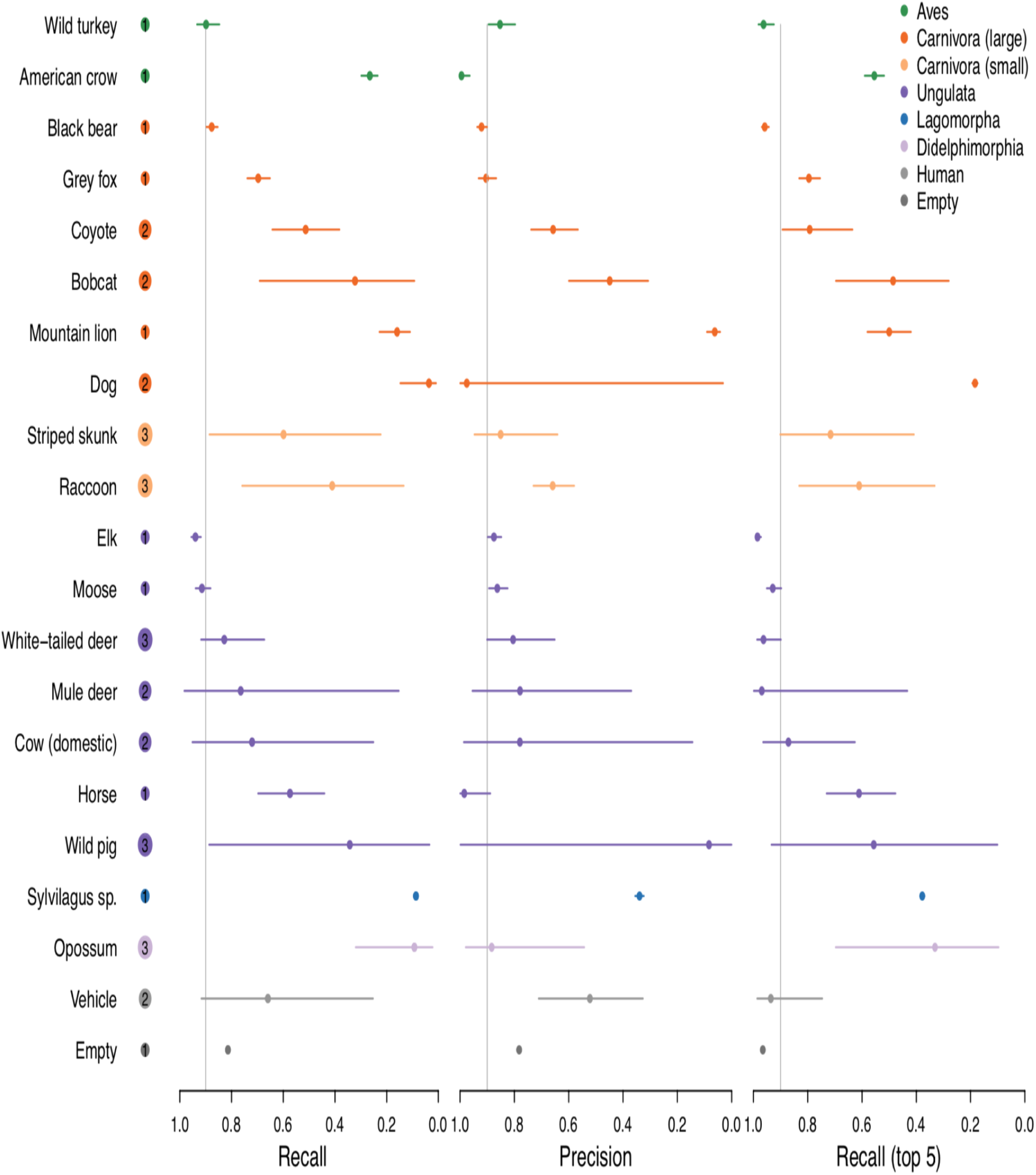
Species model out-of-sample validation revealed variable recall and precision rates across species. Median values across datasets are presented along with 95% confidence intervals. The number of datasets for each species is included in the circle next to the species name.

## 4 Discussion

In mlwic2, we provide two trained machine learning models, one classifying species and another distinguishing between images with animals and those that are empty, with 97% accuracy, which can potentially be used to rapidly classify camera trap images from many locations. While the species model performed well on out-of-sample images from Saskatchewan, Canada (91% overall accuracy), the model performed poorly on some out-of-sample datasets (Table 2; Fig. 2). The discrepancy in model performance on images from different datasets indicates that transferability remains an issue and our species model will not be useful on all datasets; some users will need to train new models on images from their field sites, an option that is available in mlwic2. Nevertheless, even in the Missouri dataset where our model performed worst, the top-5 accuracy, the rate at which the true species in an image was in the model’s top-5 guesses, was 65% (Table 2). For some applications, e.g. detection of invasive or rare species, a good out-of-sample top-5 recall rate may be sufficient to address scientific questions or meet monitoring objectives. Additionally, our empty-animal model performed well at distinguishing empty images from those containing animals in datasets from three different countries (91-94% accuracy), indicating that this model may be broadly applicable for removing empty images from datasets globally. We propose a workflow for how users can apply these models to filter-out empty images and train new models as necessary (Fig. 3). By providing Shiny apps to train models and classify images, we make this technology accessible to more scientists with minimal programming experience. Our finding that high recall (¿95%) can be achieved with fewer than 2,000 images for some species (Table 1; Fig.1) suggests that smaller labeled image datasets can potentially be used to train models with this software.

**Figure 3:**
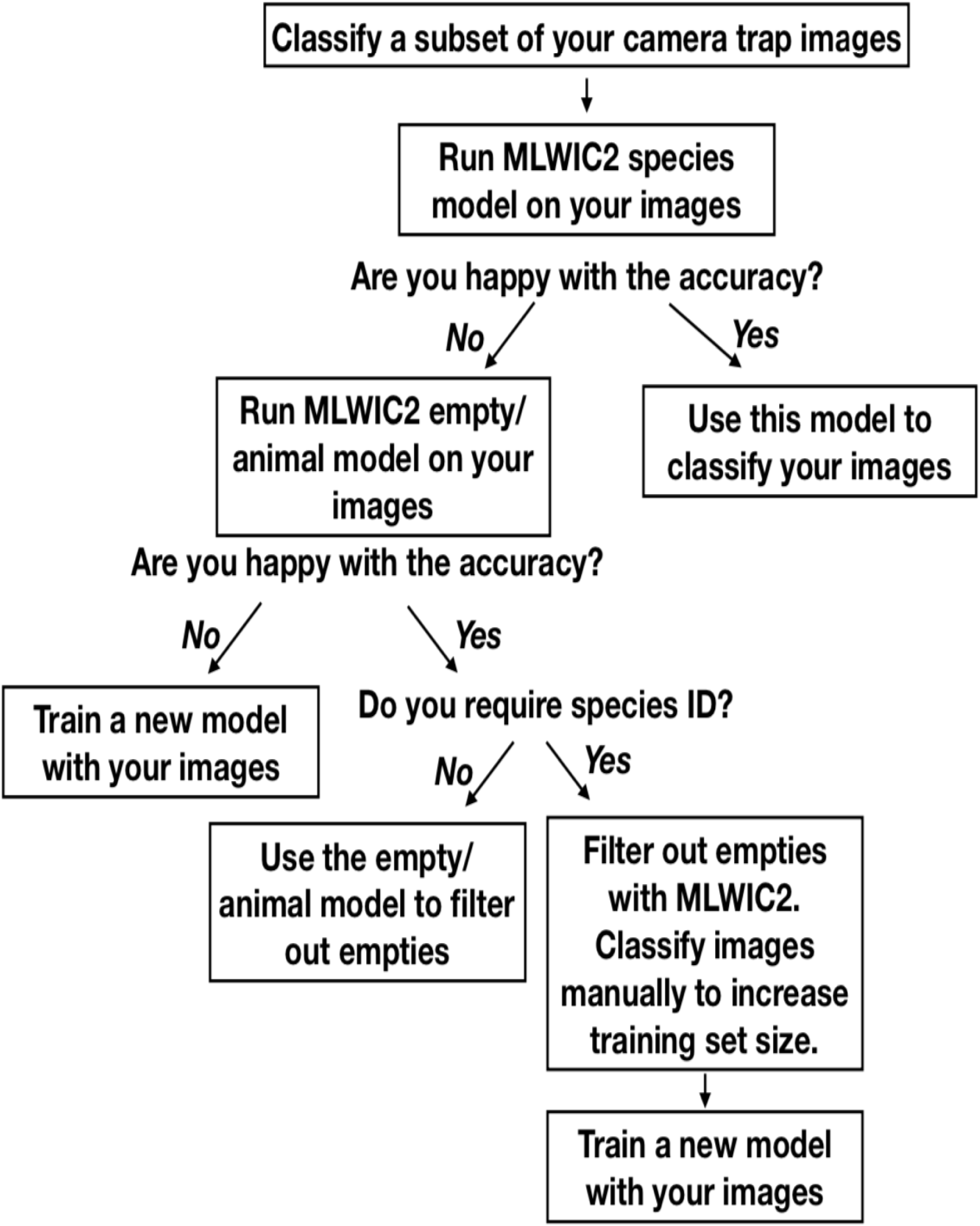
Proposed workflow for using mlwic2 models when classifying camera trap images.

Other researchers have developed models for recognizing animals in camera traps, with some success in out-of-sample identification. For example Zilong software accurately removed 85% of empty images (Wei et al., 2020), MegaDetector had a precision of 89-99% at detecting animals (Beery et al., 2019), and mlwic achieved an accuracy of 82% at out-of-sample species classification (Tabak et al., 2018, 2019). We hypothesize that our models performed well on some out-of-sample datasets (Snapshot Serengeti, Snapshot Karoo, Wellington, and Saskatchewan; Table 2) because they were trained using camera trap images from multiple locations with different camera placement protocols, allowing the model to develop a search image for each species in multiple backgrounds.

Transferability of machine learning models remains a complication for implementing these models more broadly to camera trap data and, in many cases, it is most productive for scientists to build models that are trained directly on their study sites (see Fig. 3). While such models will have less broad applicability (they are unlikely to be accurate globally), they can have high study-specific accuracies, thus reducing the burden of manual image classification.

## 5 Acknowledgements

Contributions of JCB were partially supported by the DOE under Award Number DE-EM0004391 to the University of Georgia Research Foundation. Support for this research was provided by the USFWS Pittman-Robertson Wildlife Restoration Program and Wisconsin Department of Natural Resources.

For supplying camera trap images, we thank USDA Forest Service: Rocky Mountain Research station; Montana Fish, Wildlife and Parks; Wyoming Game and Fish Department; Washington Department of Fish and Wildlife; Idaho Department of Fish and Game; Woodland Park Zoo.

## Disclaimer

This manuscript was prepared as an account of work sponsored by an agency of the United States Government. Neither the United States Government nor any agency thereof, nor any of their employees, makes any warranty, express or implied, or assumes any legal liability or responsibility for the accuracy, completeness, or usefulness of any information disclosed, or represents that its use not infringe privately owned rights. Reference herein to any specific commercial product, process, or service by trade name, trademark, manufacturer, or otherwise does not constitute or imply its endorsement, recommendation, or favoring by the United States Government or any agency thereof. The views and opinions of the authors expressed herein do not necessarily state or reflect those of the United States Government or any agency thereof. Any use of trade, firm, or product names is for descriptive purposes only and does not imply endorsement by the U.S. Government.

## 6 Author contributions

MAT, RSM, and RKBoughton conceived of the project. DWW, RKB, JSI, EAO, ESN, RYC, JLS, FI, JE, RKB, AJD, JSS, DPW, JCB, and KCV oversaw the data collection and labeling processes. MSN and JC provided insight for model training. MAT developed mlwic2 and led the writing of the manuscript. DWW and EJN assisted with mlwic2 development. All authors contributed critically to drafts and gave final approval for submission.

## 7 Data availability

The trained models described in this work are available in the mlwic2 package (https://github.com/mikeyEcology/MLWIC2). Images used to train models are available in the North American Camera Trap Images dataset (lila.sciece/datasets/nacti).

## 9 Supporting Information

Appendix S1: Information for each of the 18 studies that produced camera trap images used in this paper. The final 59 columns are the number of images of each species (or group of species).

Appendix S2: Learning rate and weight decay for each epoch in the model training process.

Appendix S3: Calculation of pooled recall and precision rate and corresponding confidence intervals.

Appendix S4: Confusion matrix depicting the number of images of each species (or group of species) that were classified by the species model as each species (or group of species). Columns are the ground truth labels from human observers; rows are predictions from the model.

